# Deep Learning on MRI Affirms the Prominence of the Hippocampal Formation in Alzheimer’s Disease Classification

**DOI:** 10.1101/456277

**Authors:** Xinyang Feng, Jie Yang, Zachary C. Lipton, Scott A. Small, Frank A. Provenzano, Alzheimer’s Disease Neuroimaging Initiative

## Abstract

Deep learning techniques on MRI scans have demonstrated great potential to improve the diagnosis of neurological diseases. Here, we investigate the application of 3D deep convolutional neural networks (CNNs) for classifying Alzheimer’s disease (AD) based on structural MRI data. In particular, we take on two challenges that are under-explored in the literature on deep learning for neuroimaging. First deep neural networks typically require large-scale data that is not always available in medical studies. Therefore, we explore the use of including longitudinal scans in classification studies, greatly increasing the amount of data for training and improving the generalization performance of our classifiers. Moreover, previous studies applying deep learning to classifying Alzheimer’s disease from neuroimaging have typically addressed classification based on whole brain volumes but stopped short of performing in-depth regional analyses to localize the most predictive areas. Additionally, we show a deep net trained to distinguish between AD and cognitively normal subjects can be applied to classify mild cognitive impairment patients, with classification scores aligning empirically with the likelihood of progression to AD. Our initial results demonstrate both that we can classify AD with an area under the receiver operator characteristic curve (AUROC) of .990 and that we can predict conversion to AD among patients in the MCI subgroup with an AURUC of 0.787. We then localize the predictive regions, by performing both saliency-based interpretation and rigorous slice and lobar level ablation studies. Interestingly, our regional analyses identified the hippocampal formation, including the entorhinal cortex, to be the most predictive region for our models. This finding adds evidence that the hippocampal formation is an anatomical seat of AD and a prominent feature in its diagnosis. Together, the results of this study further demonstrate the potential of deep learning to impact AD classification and to identify AD’s structural neuroimaging signatures. The proposed classification and regional analyses methods constitute a general framework that can easily be applied to other disorders and imaging modalities.

Acronyms

MRI: magnetic resonance imaging
AD: Alzheimer’s disease
CN: cognitively normal
MCI: mild cognitive impairment
MCIs: mild cognitive impairment stable
MCIp: mild cognitive impairment progression
HP: hippocampal formation
ADNI: Alzheimer’s Disease Neuroimaging Initiative
HCP: Human Connectome Project
CNNs: convolutional neural networks
CAM: class activation mapping
ReLU: rectified linear unit
BN: batch normalization
AUC: area under the curve
ROC: receiver operating characteristic
AUROC: area under the receiver operating characteristic curve

## Introduction

Alzheimer’s disease is a progressive neurodegenerative disease responsible for the majority of cases of dementia [Alzheimer’s Association, 2018]. Because of the degenerative nature of the illness and the current lack of a cure, much research focuses on developing techniques for early diagnosis and intervention [Nestor et al., 2004, Barnett et al., 2014]. Given an accurate early detection system, future treatments would likely have the greatest impact if administered earlier in the disease progression. Our work applies deep learning to the task of detecting clinical diagnosis of Alzheimer’s disease from magnetic resonance imaging (MRI), building upon recent studies that have demonstrated the usefulness of MRI in diagnosing AD, in recognizing mild cognitive impairment (MCI) (the corresponding prodromal stage), and in categorizing biomarkers associated with neurodegeneration in AD [Jack et al., 2016].

Among different brain MRI modalities, T1w structural MRI is the most widely available and enjoys the additional benefit of being relatively standardized across scanners and protocols. Consequently, diagnosis algorithms based on T1w structural MRIs are appealing as a potential tool to assist in disease screening given the wide availability of research scans for training models, and the ubiquity of MRI scanners in the world potentially enabling the rapid deployment of learned models via software.

Following breakthroughs in computer vision, deep learning techniques have emerged as popular tools for analyzing medical images. On standard CV tasks such as classification [Krizhevsky et al., 2012], object detection [Girshick, 2015] and semantic segmentation [Long et al., 2015], deep learning algorithms based on convolutional neural networks (CNNs) [LeCun et al., 1990] have achieved undisputed dominance. For specific tasks with abundant training data—and when the training data and test data are sampled from the same distribution, these models often achieve human-level performance or better [He et al., 2015]. Moreover, due to the generality of the methods, the availability of open source code, and the wide availability of specialized computer hardware for accelerating these algorithms, they are now easily adopted by practitioners. Over the last few years, these techniques have been widely-applied in image-aided medical diagnosis. Successful applications of deep learning in medical imaging include segmenting images produced from electron microscopy [Ronneberger et al., 2015], detecting diabetic retinopathy from 2D retinal fundus photographs [Gulshan et al., 2016], and recognizing skin cancer from photographs [Esteva et al., 2017].

Learning from 3D scans, such as MRI, presents a number of additional challenges. While the number of voxels corresponding to the 3D volume representing a single patient can be large, we still have just one label per scan, raising technical questions about how to prevent overfitting. However since many brain disorders correspond to both focal and diffuse involvement, machine learning models capable of acting upon the whole volume are appealing. To that end, we explore the use of 3D convolutional neural networks for diagnosing Alzheimer’s disease, considering a variety of techniques and including some unconventional data sources in order to learn good representations without overfitting. Although 3D CNNs have been explored in the literature addressing settings as diverse as video (where time comprises the 3rd dimension) [Ji et al., 2013, Tran et al., 2015] and medical imaging [Milletari et al., 2016, Çiçek et al., 2016, Dou et al., 2016], they are relatively underutilized compared to 2D CNNs and thus best practices for deploying these models are less firmly established.

We build upon several earlier works applying 3D CNNs to AD diagnosis, which we summarize below. Note that various papers address different datasets, and different cohorts even within the same corpus, making direct comparisons of performance across papers difficult. The earliest paper in this literature to our knowledge is due to Payan and Montana [2015], who address MRI-based AD diagnosis using 2265 scans selected from the ADNI dataset. The authors report both 2-way classification accuracy for each combination of AD, CN, and MCI, and achieving 95.39% accuracy on AD versus CN and 89.47% 3-way accuracy. In this paper, they learn the weights of the convolution filters by training an auto-encoder—autoencoding consists of learning both an encoder *e*(·) and decoder *d*(·), with the goal of producing a (typically compressed) latent representation *z* = *e*(*x*) for each input *x* such that the input can be reconstructed accurately (most often measured by mean squared error), i.e., the parameters of *d* and *e* are learned to minimize the expected reconstruction error 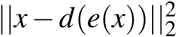. While their study provides promising support of the efficacy of 3D CNNs for diagnosing AD from brain MRI, it leaves open many modeling questions. For instance, they consider only shallow networks consisting of a single convolutional layer, followed by a pooling operation and one fully-connected layer. Moreover, they leave the autoencoder-derived filters fixed, optimizing only the weights of the fully-connected layer on the AD classification task. Following similar ideas, Hosseini-Asl et al. [2018] presented a model using unsupervised auto-encoding followed by (supervised) fine-tuning on a comparatively small dataset consisting of just 210 subjects (70 each for AD, MCI, CN) and showed impressive predictive performance with an area under the receiver operator characteristic (AUROC) of 0.993 and 99.3% accuracy in AD versus CN classification. Korolev et al. [2017] apply a 3D network architecture, achieving AUROC of .88 with an accuracy of 79% in 50 AD and 61 CN subjects. They also attempt to interpret the network, introducing a heuristic technique for feature attribution. The method consists of generating predictions while obstructing various regions in the image to determine which regions impact the model’s predictions. A recent large-scale study [Wegmayr et al., 2018] proposed using 3D CNNs directly, training weights from scratch (no unsupervised pre-training) and achieved 86% accuracy in AD/CN classification in a merged ADNI+AIBL dataset consisting of 6618 scans from CN subjects and 4476 scans from AD subjects. These studies demonstrate the promise of modern CNN architectures for extracting patterns from brain MRI.

Despite generating accurate predictions, deep learning has long been described as a black-box. In attempts to *interpret* or *explain* the classifications produced by various deep learning techniques, a wave of papers have proposed diverse heuristics for explaining predictions, including generating textual explanations [Hendricks et al., 2018], and providing feature-wise attributions [Ribeiro et al., 2016, Zhou et al., 2016], sometimes in the form of visualizations [Simonyan et al., 2014, Sundararajan et al., 2017]. The topic is intensely debated and researched in connection with critical settings like medical diagnosis, predictive policing, and other impactful automated decision-making scenarios where accountability is a concern [Caruana et al., 2015]. A recent study [Yang et al., 2018] utilized multiple methods to generate visual explanations for AD classification despite relatively low classification performance (0.863 AUROC, 0.766 accuracy) ona small dataset consisting of 47 AD and 56 CN subjects. However, despite intense interest in the broader topics of interpretability/explainability, Lipton [2018] showed that the very definition of these concepts remains elusive. For many interpretability techniques, the papers fail to clarify precisely what task they are addressing, let alone whether they are successful. Ablation tests constitute one classic method for probing a predictive problem to assess the usefulness of various features. These tests consist of assessing predictive performance when individual features or sets of related features are dropped from consideration. Our study complements the more speculative modern interpretability analyses with rigorous ablation tests to assess the predictiveness of each region of AD.

In this study, we build upon previous work applying 3D deep convolutional neural networks to AD diagnosis, using a large-scale structural MRI dataset. In particular, we focus on two specific aspects: (i) incorporating longitudinal scans as a unconventional data source, and (ii) an investigation aimed at localizing the most predictive regions. Generally, data augmentation helps to prevent models from overfitting by enriching an original source of data through the addition of variations of these examples perturbed through transformations with respect to which we would like the model to be invariant. In typical photographic images, such transformations might include random crops, translations, rotations, and small changes to the color palette [Krizhevsky et al., 2012]. Here, instead, we enrich the data by including images captured from the same patient across multiple visits. Inclusion of multiple scans from the same subject raises two important issues: the data leakage problem and the disease progression problem. Data leakage occurs when the training and test sets contain the different scans from the same subject, the model might make the prediction by memorizing and retrieving the label from the same person and is likely to result in over-optimistic performance. This issue is dealt with easily by ensuring that the partition into train/validation/test splits takes place at the level of the individual patients instead of at the level of the scan. The disease progression problem relates to the fact that the disease status of subjects might change during follow-ups, and the cross-sectional diagnosis labels for a certain scan might be different from the baseline label, this is especially important in prodromal disease status, e.g. MCI in AD pathology. We then might reasonably ask how many of these patients have truly undergone a change in status (versus being initially mislabeled). Note that while including subsequent scans raises this question, it does not cause the problem of mislabeled baseline scans.

Most previous studies on AD classification with CNNs [Hosseini-Asl et al., 2018, Korolev et al., 2017] exploit the (single) baseline scan for each subject and train their models to predict the cross-sectional diagnosis labels (assessed at scan time). Payan and Montana [2015] utilizes multiple scans but did not explicitly address potential data leakage and disease progression problems. Wegmayr et al. [2018] explicitly addresses the data leakage problem (they refer to it as *subject duplication*), ensuring non-overlapping subjects in training and test sets, however, the disease progression is not explicitly discussed while stating several scans of an individual subject typically have the same disease label, which could be potentially problematic especially since they performed a three-way classification including MCI. Another difference from this previous study is that we opted to only include multiple scans from different sessions rather than within-session duplicate scans to leverage the data richness versus data redundancy.

Our model achieves high classification performance in AD classification in a large dataset consisting of 2817 scans/sessions from CN subjects and 1874 scans/sessions from AD subjects. Moreover, we find that by applying model trained on AD versus CN classification to baseline scans of MCI subjects, we can accurately predict progression from MCI to AD. Furthermore, we study the neuroanatomical underpinning in AD classification with a series of novel regional analyses following the regional vulnerability idea [Small, 2014]. We pinpoint hippocampal formation as a most predictive driver for our deep-learning-based AD diagnosis model, which further affirms the prominence of hippocampal formation in AD classification.

We explore regional significance in a number of ways. In one approach, we generate a 3D activation map *post hoc* without any changes to the model or input data, presenting the whole brain volume to the classifier and inferring those regions that contribute most to the classification using a 3D class activation mapping (CAM) technique [Zhou et al., 2016]. CAM utilizes fully convolutional neural networks whose output is produced by a global pooling operation followed by the softmax operator to generate the predicted probabilities. By weighting the pre-pooled activation maps with the corresponding input to the softmax operator, CAM can assess per-region importance to the classification, generating a visual map indicating which voxels are suggestive of the target label. Gradient-weighted CAM (grad-CAM) [Selvaraju et al., 2017] generalize the CAM to broader CNN families by flowing the gradients of the target label into the last convolutional layer. In our case, we utilize grad-CAM to determine which specific 3D regions most indicate a prediction of AD.

We also explore two “ablation”-based methods that explicitly focus on specific brain regions. The first method consists of training models to predict AD using 2D MRI slices (in each of the three coordinate planes), and evaluate the pattern of the capabilities of slices differentiating AD. The second method, more informed of neuroanatomy, consists of masking specific regions using masks generated from a sub-population with segmentation, as described in detail in Data and experimental setup, training on the masked regions, and comparing the classification accuracy.

The CAM method reveals a preponderance of activation overlap containing the left anterior hippocampal formation (HF). Evaluation on 2D MRI slices demonstrates the importance of slices covering the HF in the classification of AD using deep 2D CNN model. And evaluation on isolated brain lobes also demonstrates the importance of temporal lobe, which contains the HF. These findings have implications for both the interpretability of CNNs used in image-based disease diagnosis and also the prospective MRI acquisition protocols targeting AD diagnosis. Even for highly complex and nonlinear models, regionality and the underlying pathology revealed still manifests importance.

Importantly, we note that our proposed 3D CNN model and regional analyses constitute a highly general framework that can potentially be applied to other brain disorders and imaging modalities.

## Methods

### Data and experimental setup

The dataset used in this study is from the Alzheimer’s Disease Neuroimaging Initiative (ADNI) ^1^. The details about the MRI data acquisition can be found in ADNI website ^2^. The T1-weighted structural MRI scans were pre-processed with the standard Mayo Clinic pipeline ^3^. The AD diagnosis was based on clinical evaluations. The MRI and diagnosis data were queried and accessed at August 2017. The diagnosis includes AD, MCI, and cognitively normal (CN). The dataset used to generate the regional masks includes 382 scans from unique elderly subjects in ADNI-2 and 1113 scans from unique young subjects in Human Connectome Project (HCP) ^4^ [Van Essen et al., 2013]. The subjects cover age 25-90, and clinical diagnosis of young normal control (*N* = 1113), elderly normal control (*N* = 120), MCI (*N* = 138) and AD (*N* = 124).

For the experiment of AD versus CN classification, we sought to include as many MRI sessions as possible that correspond to an AD or CN diagnosis at scan time. Specifically, we included the baseline and follow-up scans of patients diagnosed as AD at baseline, the baseline and follow-up scans of subjects diagnosed as cognitively normal at baseline before the conversion to AD or MCI if ever happened, and the after-conversion follow-up scans of subjects who were CN or MCI at baseline but later progressed to AD. In total, we include 4691 scans (2817 CN, 1874 AD) from 1189 subjects under these criteria. This experiment setup basically sets an upper limit to the amount of data for cross-sectional AD versus CN classification in ADNI cohort. The inclusion of scans after conversion also helps enrich the samples around the “classification boundary” as will be more thoroughly discussed in the “Inclusion of longitudinal scans” section.

We used 8/10 of the subjects as training set consisted of 3709 scans (2240 CN, 1469 AD) from 952 subjects, 1/10 as validation set consisted of 456 scans (279 CN, 177 AD) from 119 subjects, and 1/10 as test set consisted of 526 scans (298 CN, 228 AD) from 118 subjects. As discussed in the Introduction section, the training, validation and test sets split was partitioned at subject level through stratified random sampling on baseline diagnosis labels so that the groups have non-overlapping subjects and approximately even distribution of baseline diagnosis labels. The final model is selected as the one that has the highest validation accuracy (i.e. classification accuracy in validation set).

For the experiment of MCI progression prediction on baseline scans, we trained another model only using subjects whose *baseline* diagnosis are cognitively normal or AD to prevent data leakage. We included scans from 796 subjects under this criterion. Similarly, 2918 scans (1943 CN, 975 AD) from 626 subjects were used as training set, 382 scans (251 CN, 131 AD) from 80 subjects were used as validation set, 325 scans 229 CN, 96 AD) from 80 subjects were used as test set. We used the same neural network training setup.

We included 318 MCI stable subjects and 311 MCI progression subjects for MCI progression prediction. The MCI stable subject are those who remained MCI during a follow-up period of at least 3 years from the initial visit. The MCI progression subjects progressed to AD at follow-up visits, among which 256 subjects progressed to AD within 3 years.

We used the area under the curve receiver operating characteristic (AUROC) and accuracy to evaluate the classification performance.

### Image preprocessing

Basic pre-processing steps include nonparametric nonuniform intensity normalization (N3) based bias field correction [Sled et al., 1998], brain extraction using FreeSurfer [Ségonne et al., 2004], and 12 degree of freedom affine registration (using FSL FLIRT [Jenkinson et al., 2002] with normalized mutual information cost function) to the 1*mm*^3^ isotropic MNI152 brain template. The dimension of the 3D volume is 182 × 218 × 182 (LR × AP × SI).

Bias field correction is generally robust, fast, and based on physics models which act as a strong prior [Sled et al., 1998]. There are brain extraction methods using deep learning techniques, but there is not one that is well-validated and widely-available. Skull-stripping using FreeSurfer in general provides consistently high quality brain extraction.

The registration is to ensure same orientation and roughly same spatial correspondence of different images. Although there are techniques such as spatial transformer network [Jaderberg et al., 2015] to learn transformation within the network, it would involve more parameters to learn and add burden to the data.

All the brain extracted, affine-transformed images were checked by a well-trained reviewer with visual inspection. Scans having severe MRI artifacts, brain extraction failure or poor registration were excluded.

### Inclusion of longitudinal scans

Commonly, computer vision practitioners synthetically augment their datasets, applying random transformations to the existing training images, including translation, rotation, scaling, etc. However, unlike natural images or those collected from some other medical imaging modalities, where objects of interest might vary in location and rotational orientation, MRI images of brains are approximately at the same position through registration, with the brain regions roughly aligned. Thus in this setting, learning rotational and translational invariances is not well motivated.

There is another form of data augmentation or more precisely “data source” specific to medical imaging applications. For longitudinal studies, test-retest studies and just ordinary studies, there might be multiple scans per subject. By including time as a factor in subject identification, we can increase the amount of data. In a sense, by including these data sources, we are seeking natural forms of data augmentation. The corresponding “transformations” in data augmentation would be normal aging or disease progression or both (longitudinal scans with a significant interval between scans), subject re-positioning (scans acquired at different sessions and within a short period of time) and subject motion (scans acquired at the same session). The variability present in the scans or the data coverage in the whole data space decreases in this order. The illustration of the general idea is shown in Figure 1. As discussed in Introduction section about disease progression, special attention is required for the first kind, where the different time points of the same individual might be at different health or disease stages. Moreover, those scans, lying on the verge of different diagnosis, constitute informative cases for the classification.

**Figure 1.**
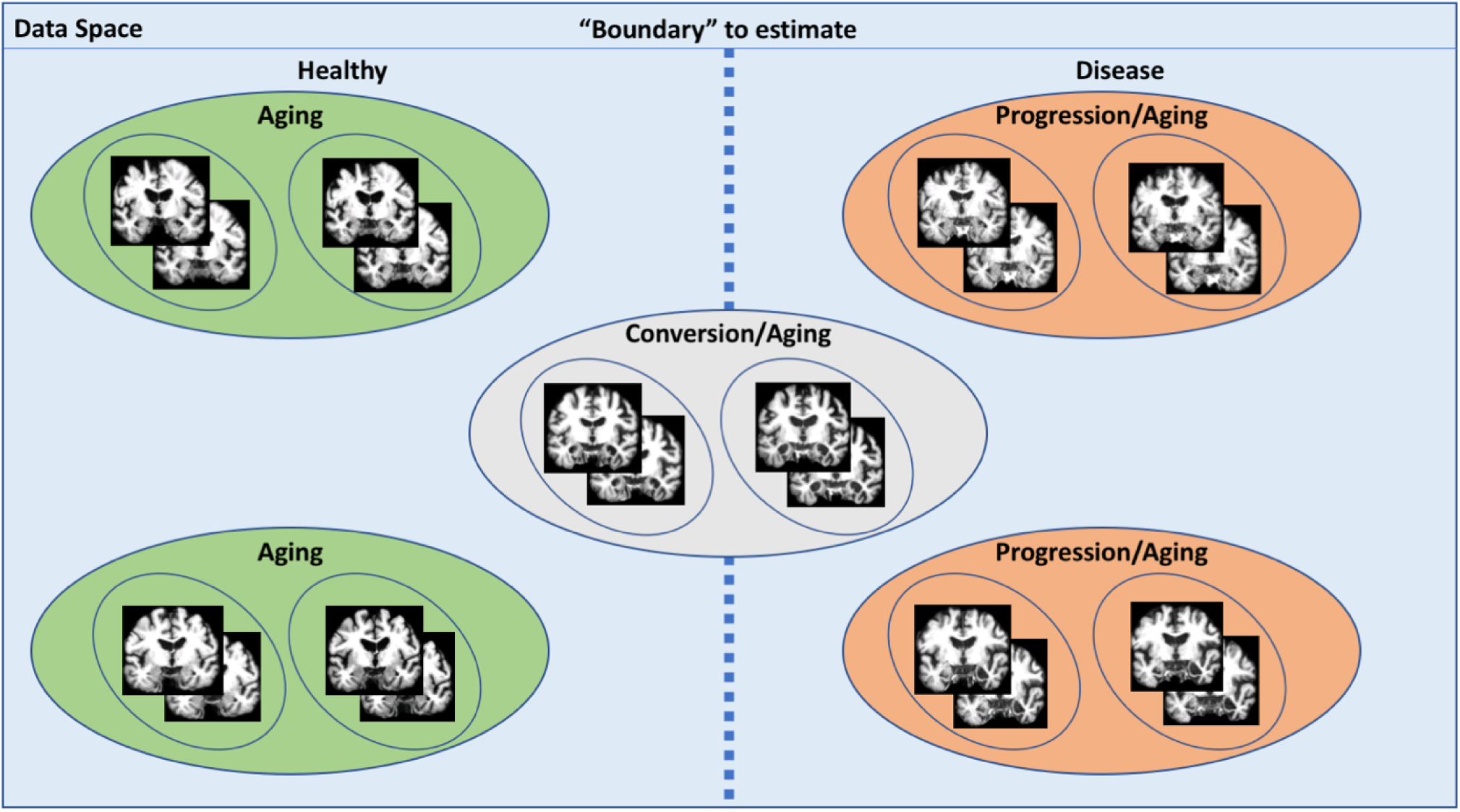
Illustration of data augmentation or inclusion of longitudinal scans in MRI studies. The whole plane is a simplified representation of the data space. Each large circle indicates one individual subject, each small circle indicates one MRI session. Each coronal slice of MRI scan represents one scan. The objective of the deep learning algorithm is to find the “boundary” (dashed line) that best differentiates cognitively normal subjects and AD patients. Enriching our data by using longitudinal scans from subjects helps to increase the data coverage from the small circle to the large circle.

In this study, we opt for using scans from different sessions, which already provides a significant increase in the amount of data: from 796 baseline scans to 4691 scans. And scans from the same scanning session present very low variability.

### Convolutional neural networks

We use a general CNN architecture similar to the VGG classification architecture [Simonyan and Zisserman, 2015] with multiple interleaved convolutional blocks and max pooling layers and increasing number of features along the depth. The main differences include having just one fully-connected layer to significantly reduce the number of parameters and replacing 2D operations with 3D operations. For convolutional layers, we use a convolutional kernel size of 3 × 3 × 3, a batch size of 5, rectified linear unit (ReLU) as the activation functions. We flatten the output from the last convolutional layer and feed into a fully-connected (FC) layer with sigmoid as the activation. We use batch normalization (BN) before the activation function. The algorithm was optimized using Adam method with cross-entropy loss function. The initial learning rate was tuned in the range from 1*e* − 4 to 1*e* − 6 including [1e-4, 5e-5, 2e-5, 1e-5, 5e-6, 2e-6, 1e-6] and was set at 2e-5. We implemented the algorithm using Keras and TensorFlow. As early stopping criteria, we set the patience parameter on validation accuracy to 10 epochs. We include weight *l2* regularization (also known as weight decay) to prevent overfitting with a factor of 1.0. An illustration of the framework is shown in Figure 2. In this study we use five (N in Figure 2) stages. The feature dimension of the first layer is 16 and increases by a factor of 2 in each subsequent stage.

**Figure 2.**
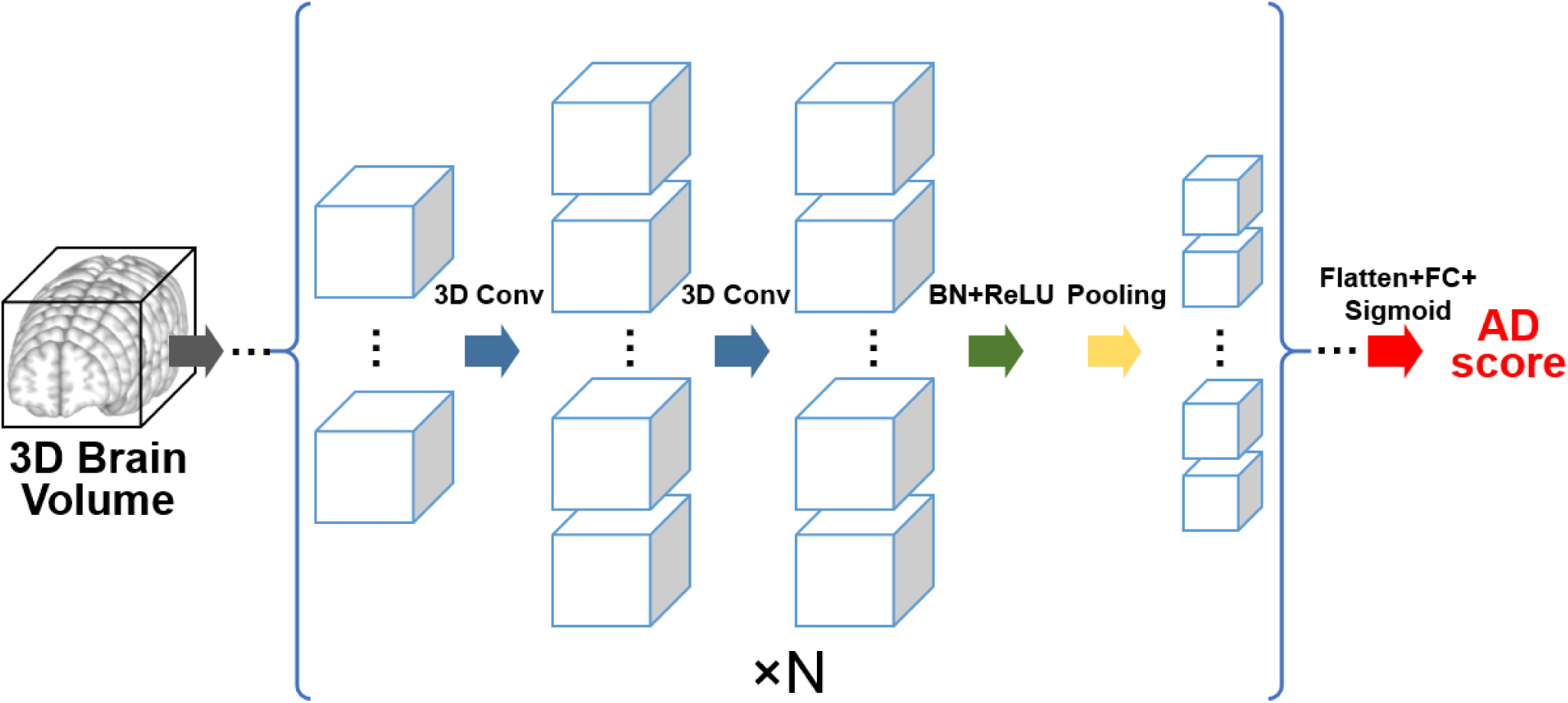
The convolution neural network architecture. The inputs are 3D brain volumes. Each cubic represents one 3D feature map, the size reflects the spatial dimension of the feature map, the number reflects the number of feature maps (channel dimension). The blue arrows are 3D convolutional operations, the green arrow represents batch normalization (BN) followed by rectified linear unit (ReLU), the yellow arrow denotes the max pooling operation. The basic unit enclosed in the bracket is repeated *N* = 5 times with increasing number of features and decreasing spatial dimension. The final convolutional output is flattened and fed into one fully-connected (FC) layer with sigmoid activation function (red arrow), generating the final AD score.

### Application to MCI progression prediction

The classification was trained on AD versus CN, which presents the largest neuroanatomical contrast in the AD spectrum. Whether the model learned using the two ends of the spectrum could inform the differentiation of patients in the middle of the spectrum is critical. We directly applied the model to the baseline scans of patients diagnosed with MCI since MCI patients are not part of the training dataset of the model. We use AUROC to evaluate the prediction performance.

### Class activation map

Class activation map (CAM) original proposed by Zhou et al. [2016], extended and generalized in Gradient-weighted CAM (grad-CAM) Selvaraju et al. [2017], has been used in medical image analysis field to inform the “attention” of the 2D classification [Feng et al., 2017]. In this study, we generated 3D class activation maps to visualize the predictive contribution of brain regions to the AD classification task. Specifically, we used grad-CAM with ReLU gradient modifier and rescaled generated CAM with min-max normalization. Importantly, since the map can be generated individually, it has the potential to be used as an individual neuroanatomical validity report without sacrificing the prediction power of whole brain based prediction model. We generated the average class activation map for all AD patients to demonstrate the average “attention” of the algorithm.

### MRI 2D slice based classification

Besides the *post hoc* saliency map based class activation map method, we also propose ablation analyses methods focusing on part of the input data, We tested the classification using 2D CNN with the input being three consecutive slices as three channels. This design takes the inter-subject alignment precision into consideration (i.e. not extracting just one slice) and also ensuring relative similarity among different channels (i.e. not extracting five slices). The network architecture is the same as the 3D CNN architecture described above except that the 3D operations are all replaced with the corresponding 2D operations. We report the classification performance on the different groups of 2D slices as the indication of predictive importance.

### Brain lobe based classification

Slice-based regional analysis method provides a way to investigate the predictive regions of the classification from imaging perspective, as the coordinate planes are imaging planes. But each slice still represents a mixture of multiple regions located at a certain spatial level. It is more appealing to generate neuroanatomically meaningful regions and perform classification focusing on these regions separately. A probabilistic spatial distribution of different regions was derived from the affinely co-registered FreeSurfer segmentations [Fischl et al., 2002, 2004] from 1,495 scans as detailed in the Data and experimental setup. An occurrence probability of 0.5% was used as the threshold for the lobe mask generation. The definition of lobes in FreeSurfer segmentation nomenclature is referenced in FreeSurfer website ^5^. We focused on the lobe-level ablation study firstly because the brain lobes are functionally and structurally distinct units, and also because finer region parcellation results in poor overlap across subjects and inevitable involvement of neighboring regions.

## Results

### AD classification

In Figure 3, we show the classification performance in AD versus CN task. We evaluate our model both on unique MRI sessions and the baseline scans of unique subjects (see Methods section for more details). Our model achieves 0.980 AUROC and 93.3% accuracy when evaluated on unique MRI sessions, and 0.990 AUROC and 96.6% accuracy when evaluated on the baseline scans of unique subjects. The high overall classification accuracy of our model lays a solid foundation for our subsequent results investigating regional attribution. The model training process with only baseline scans under identical training settings stuck at uniform predicted label.

**Figure 3.**
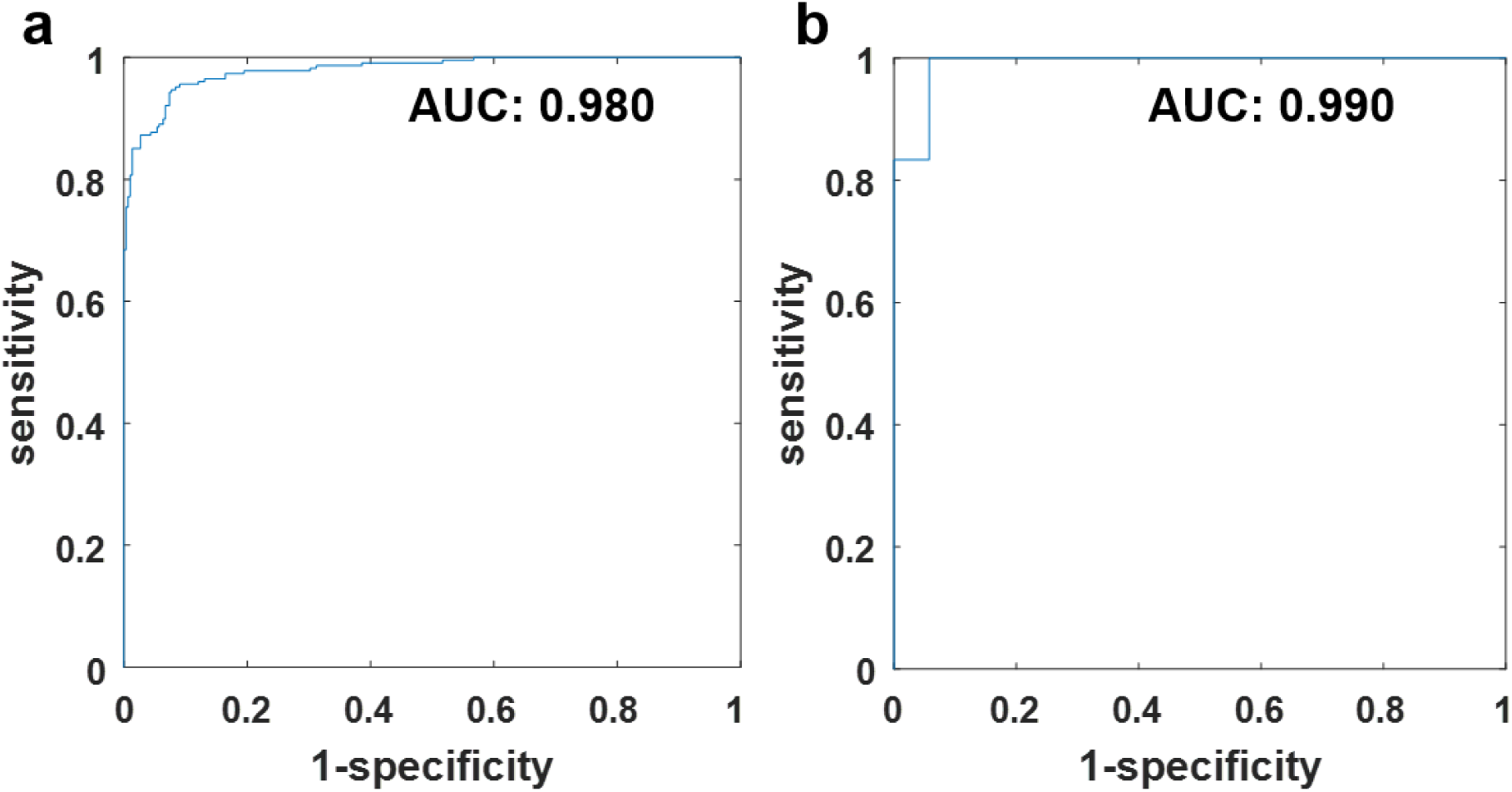
ROC curve for AD classification on the test set of a) all unique MRI sessions, b) baseline MRI scans of unique subjects. The AUROCs are annotated in the figures.

### Application to MCI progression prediction

Remarkably, we find that the classifier trained exclusively on AD and CN patients can also be used *post hoc* to differentiate among those MCI patients who will and will not progress in the near term to AD. The ADNI dataset contains MCI patients whose subsequent progression or not to AD has been noted longitudinally.Ideally, we might train a model exclusively on MCI patients whose subsequent progression status has been observed, directly learning to distinguish AD’s prodromal stage from other causes of MCI. However, the ADNI dataset does not contain sufficient MCI patients (around 600) to train such a classifier. Although the subset of MCI patients is too small for direct training, it is sufficiently large to serve as an evaluation set.

To determine the usefulness of our AD vs. CN classifier for recognizing those MCI cases that will progress to AD, we fed MCI patients through an AD vs. CN binary classifier, interpreting a higher probability of AD as more likely to progress to AD and a higher probability of CN as less likely to progress. For this experiment, we trained the AD vs. CN model *using only baseline scans from subjects diagnosed as either AD or CN at baseline* (as detailed in Data and experimental setup section) achieving an AUROC of 0.973 on i.i.d. holdout data. Then we fed our evaluation set of MCI patients through the classifier, achieving an AUROC of 0.787 (0.808 when including only MCI patients who progressed or stayed stable within 3 years), matching state-of-the-art performance while using structural MRI data only [Korolev et al., 2016]. In Figure 4, we plot the performance of our classifier as applied *post hoc* to the task of predicting MCI progression. Note that this evaluation procedure applies the CNN out-of-sample to a subset of patients that are not represented in the training set. In general, machine learning are liable to break under distribution shift and thus our performance, despite matching the previous state-of-the-art, might be far from the ceiling of what we might achieve given adequate data. Likely, in the future, given a large dataset of MCI patients, sufficient for training a progression prediction model directly, we might achieve significantly higher predictive accuracy. This could potentially imply that the neuroanatomical pattern of MCI partially lies on the normal-to-AD continuum.

**Figure 4.**
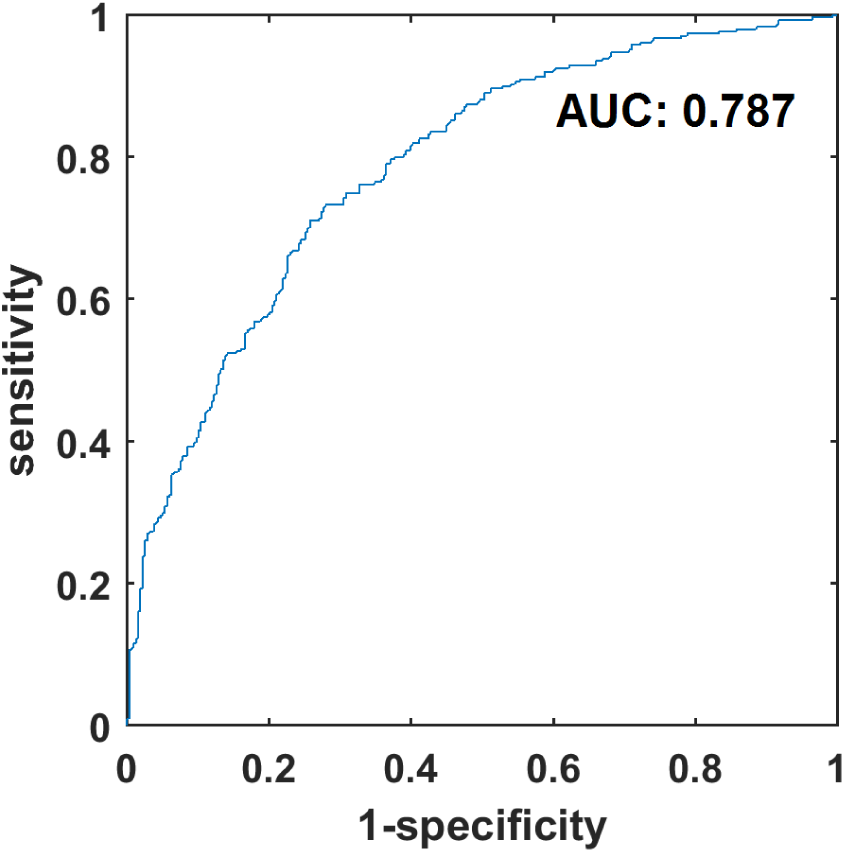
ROC curve for MCI progression prediction. The AUROC is annotated in the figure.

### AD/Class activation map

We illustrate the average class activation map for all AD patients overlaid on the MNI152 template in Figure 5. A 3D rendering of the class activation map iso-surface overlaid on the brain region segmentation provided by the FreeSurfer CVS_avg35 in MNI space is shown in Figure S1 in the supplementary material. We can see from Figure 5 and Figure S1 that the average AD/class activation map shows large “activation” in hippocampal formation, suggesting the importance of hippocampal formation in differentiating AD in our 3D deep CNN model.

**Figure 5.**
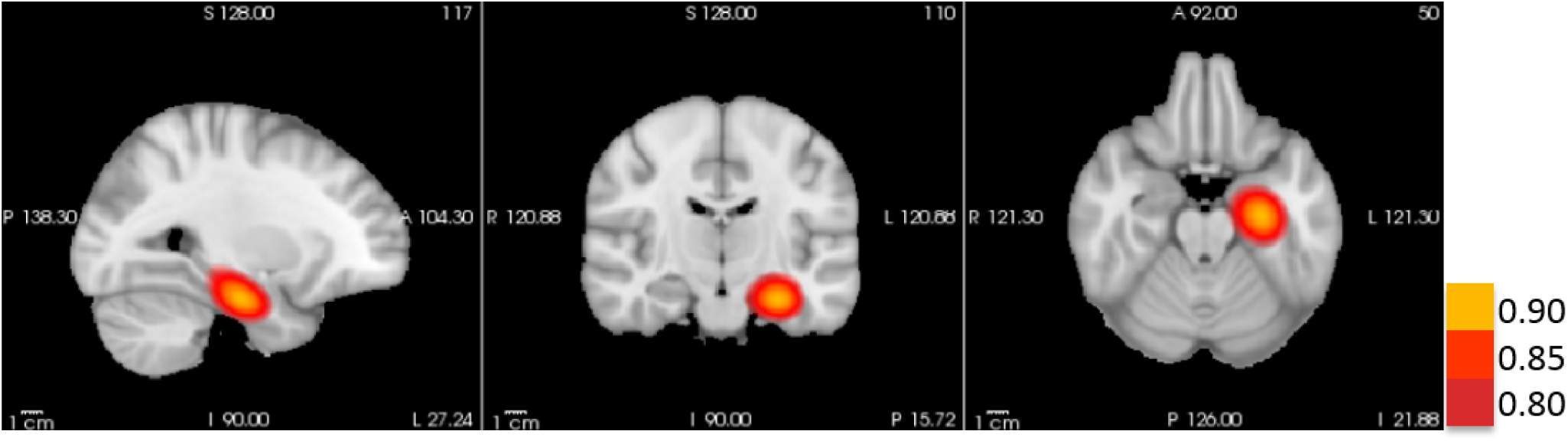
Average class activation map of AD classification overlaid on the MNI152 MRI template. The hotspot is on the hippocampal formation. The class activation map is thresholded at 0.8.

### MRI 2D slice-based classification

We also explored a slice-based classification scheme to determine which slices are most predictive of AD (Figure 6), running the analyses three times, using 2D slices along the sagittal, coronal, and axial dimensions. Along each dimension, those slices achieving highest classification performance include voxels belonging to hippocampal formation.

**Figure 6.**
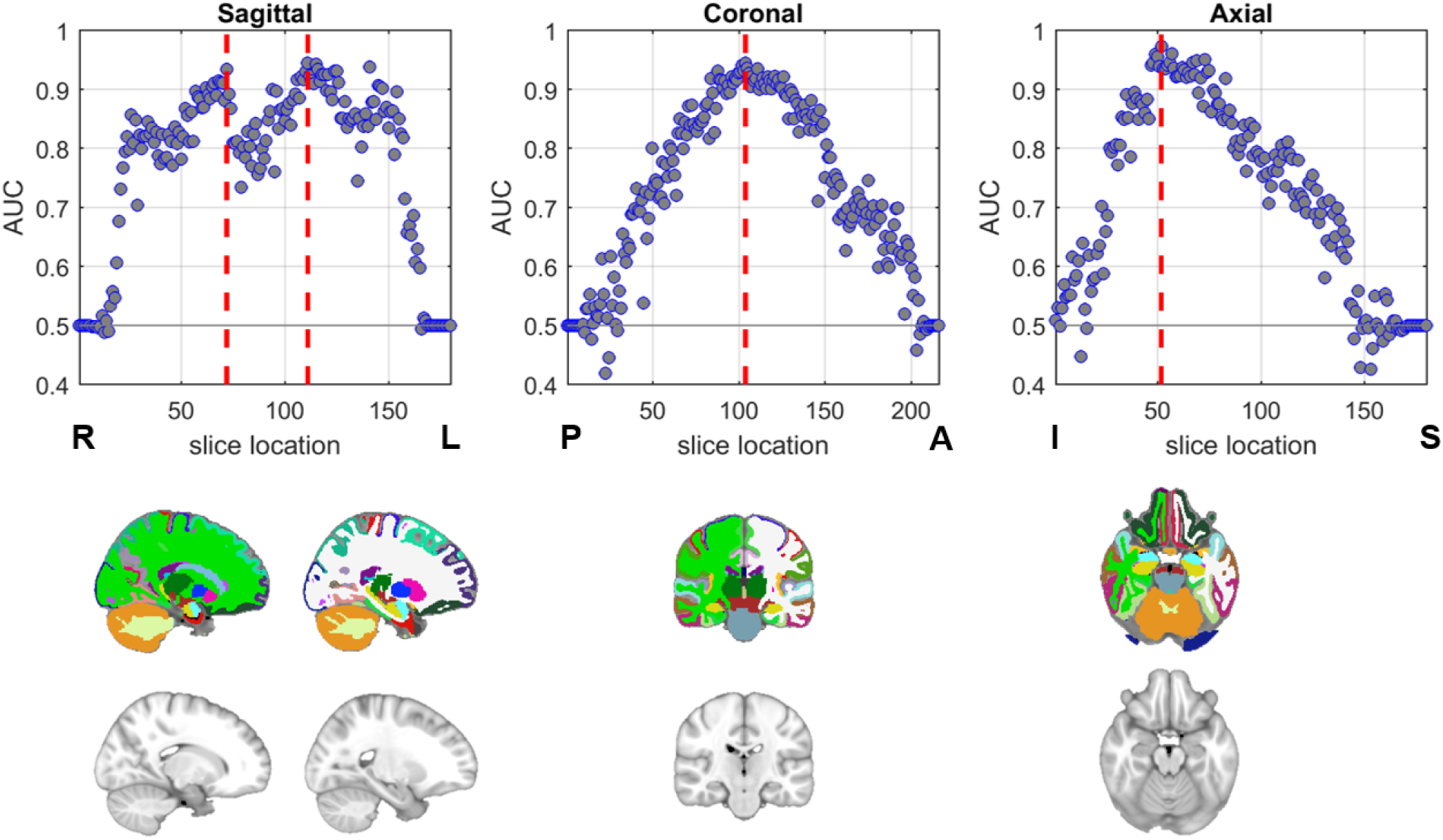
MRI 2D slice based classification. (Top row) the classification AUROC on the test set using 2D slices at different locations, the red lines indicate the location with highest AUROC. (Bottom row) the illustration of slices at the red line in the top row from the MNI152 template and the corresponding segmentation (the colors follow the FreeSurfer color lookup table: yellow-hippocampus, red-entorhinal cortex).

### Brain lobe based classification

In addition to the evidence for the importance of hippocampal formation provided by our 2D slice-based analysis, we also explored a more anatomically-informed method. Here, we explicitly train the model on different lobes and cerebellum, masking the others with the masks derived from a population probabilistic map. We illustrate these different masks in Figure S2 in the supplementary material. As shown in Figure 7, the model trained on the temporal lobe, which includes hippocampal formation, achieves the highest AUROC of 0.944 and accuracy of 88%, compared to 93% accuracy using the whole brain. The next most predictive lobe was the frontal lobe (0.899 AUROC and 83% accuracy).

**Figure 7.**
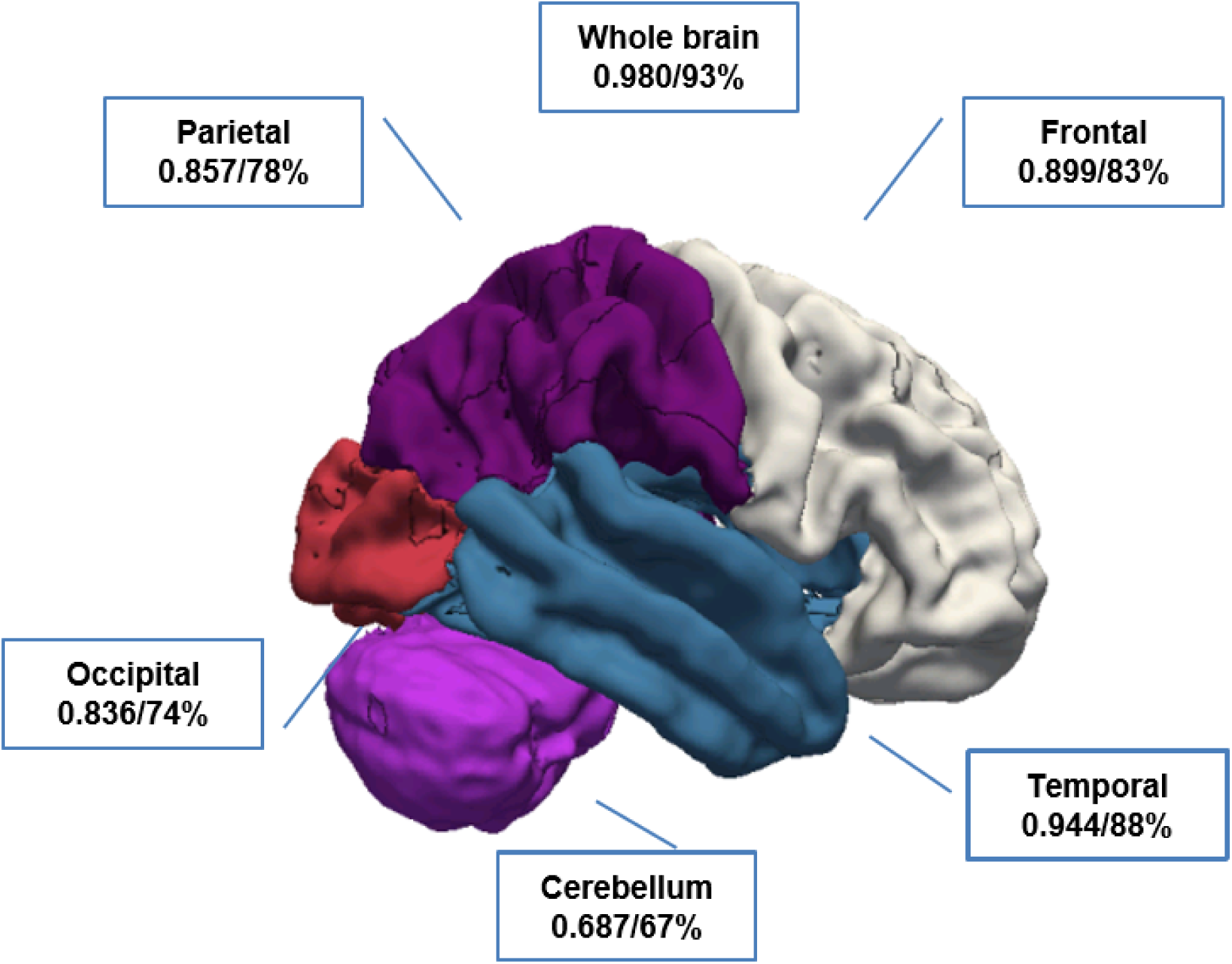
AD classification AUROC and accuracy achieved from individual lobes and cerebellum using masks from Figure S2

## Discussion

### AD staging and dysfunction spread

In Braak staging of AD [Braak and Braak, 1991], the dysfunction represented by neurofibrillary tangle starts from transentorhinal cortex (TEC) (stage I, II) to limbic regions (stages III and IV) and spreads to neocortical at stage V and VI. Additionally, a previous fMRI study has suggested cortical spread of dysfunction originating from lateral entorhinal cortex (LEC) [Khan et al., 2014]. While our findings cannot by themselves establish pathophysiological primacy, they provide evidence of structural prominence for the hippocampal formation. These results support the theory that the area circumscribing the anterior hippocampal formation is one most affected structural region in AD [Killiany et al., 2002].

### Localization

Our regional analyses show that multiple approaches of localization, including class activation maps, slice and brain lobe level ablation experiments, all suggest that the hippocampal formation is the region most predictive of AD. While the interpretation of traditional mass-univariate methods [Friston et al., 1994], via voxel-level maps may be more familiar in the medical imaging community, CNNs are often considered harder to interpret, owing to complex nonlinear patterns and interactions among voxels that these models capture. As a result, they are often considered to be black boxes, useful for pattern recognition and classification but less amenable to interpretation. The trade-offs between separability and interpretability have already been discussed in multivariate based analysis, and is becoming more obvious with the more complex architecture of deep neural networks.

However, this work highlights that through a combination of evidence produced by both heuristic saliency-based interpretations and rigorous region and lobar level ablation studies (our 2D slice-based models and lobe-masking experiments), CNNs can be used not only for predictions but also to provide insights with likely neurobiological consequence. This work presents an important case study bridging the separability and interpretability.

While the hippocampal region does appear especially predictive of AD, we emphasize that all regions offer some predictive value. Thus, in practice, for building tools to aid in the diagnosis of AD, and for predicting progression to AD among the MCI population, we recommend training models that act upon whole brain volumes. Indeed, our models acting upon whole brain volumes achieved the best AUC as compared to those acting upon any single slice or lobe.

This work relates to the multiple pathologies observations in AD [Power et al., 2018], and argues the focal neuroanatomical atrophy in hippocampal formation potentially unifies Alzheimer’s disease in the presence of multiple pathologies. Since we do not have a quantitative and meaningful local measure of structural information from structural MRI to do simple voxel-based analysis or linear multivariate analysis, we deem our current approach best suited for validating this argument.

### Prodromal disease classification using progressed cases

This study suggests the value of training a model on anchored disease or non-disease states with the utility of applying that model on classifying prodromal disease states; in this case MCI subjects who progress to AD. Many neurological and psychiatric diseases have known prodromal states, reflecting either mitigated or absent symptoms, or biomarkers that do not meet criteria for disease. These individuals are considered in a “high-risk” category, whereby only future classification (or failure to convert within a time frame) determines whether an individual had prodromal disease or another distinct illness. Neuroimaging may be sparse for these groups (due to difficulty in recruiting, for example, especially for rarer diseases) and such neuroimaging findings may be too subtle for traditional volumetric or segmentation based single subject analysis. By using a deep learning network enriched with data from confirmed disease states and controls, such a network may have value in screening for disease in broad populations, where the advanced disease has a distinct structural imaging signature which can easily and quickly be applied to high-risk states.

### Applicability

One advantage to this approach compared to many other Alzheimer’s neuroimaging findings is the range of MRI images onto which this technique can be feasibly applied. Given that the only pre-processing steps required are brain extraction and co-registration, this technique could provide a classification in a few minutes for any comparable T1-weighted image. Furthermore, the model can be retrained with more data, including data with noise or potential artifact that might otherwise prevent other standard imaging analytic techniques. It is possible that the retrained model could account for such artifacts better than traditional analytic streams.

### Practicality

The localization observations also have potential implications for acquisition. Acquisition targeting a focal field of view can facilitate diagnosis of specific disorders. And, it is important to note that the proposed method is able to generate a summary score for the structural AD-like pattern with very low computational cost at inference comparing with most of the current segmentation-based models. This is very critical in timely diagnosis and evaluation. Additionally, the computational requirement for a study this size was modest, both in cost and time, and easily adaptable into a pipeline.

### Limitations

This study has several important limitations. Firstly, each MRI, even when multiple scans from the same subject are available, are treated individually. Thus our dataset contains multiple scans from the same individual and thus those individuals that have received many scans are over-represented relative to those who undergo fewer scans. Distribution of the number of scans per subject can be found in Figure S3 in the supplementary material. In order to ensure that the multiple scans per person do not introduce any target leaks into our learning problem, we take care to construct our training, validation and testing splits on a per-individual (vs. per-scan) basis, isolating each subject to only one split. Still it is worth noting that the most heavily scanned individuals contribute more heavily to the learned model and to the evaluation. Conceivably, if one were interested in assessing the accuracy of predictions on a per-individual level, one might re-weight the scans in the test set so that each individual contributed equally to the final score.

Additionally, we note that our labels used in both training and evaluation are not ground truth *per se* as they are based on physicians’ assessments via clinical criteria *in vivo* while true diagnosis can only be confirmed at present through neuropathology, requiring post-mortem brain biopsy. We suspect that although ante- and post-mortem concordance is generally found to be high, the availability of ground truth labels would improve our models [Beach et al., 2012].

The probabilistic spatial distribution of different anatomical regions used in the brain lobe based classification experiment can be further optimized to be more reflective of general population but is not the aim and beyond the scope of this paper.

The current study aims to make predictions about the diagnosis at the time of scan, which permits different diagnosis labels of multiple scans of the same subjects. Future studies with the goal of predicting longitudinal progression may define the class labels based on the most recent diagnosis label and the follow-ups.

### Future work

Our framework is sufficiently general that it can be easily extended to other diseases such as schizophrenia, Parkinson’s disease (PD), etc. and to other MRI contrasts such as CBV, CBF or even to other imaging modalities such as PET, SPECT. One promising direction is a large-scale CBV study enabled by the recently proposed retrospective CBV technique [Feng et al., 2018]. Moreover, our ability to learn good representations for AD prediction could be brought to bear on other problems that might be supported only by smaller datasets. Following a number of successes in deep learning applications ranging from computer vision to natural language processing, we might employ transfer learning, fine-tuning the representations from our AD predictor, together with other sources of information such as age, gender, functional imaging measures, neuropsychological measures, CSF biomarkers, etc. to new tasks.

We are also interested in evaluating cross-sectional age trend, longitudinal progression, and test-retest reliability. Moreover, we would like to follow-up upon some qualitative observations observed in our localization experiments. For example, for some scans, the class activation map attributes predictions to only one side. While this interpretation technique is heuristic and does not by itself tell us anything conclusive about the localization of AD, this might suggest a hypothesis worth exploring that in some patients, at some stage in their illness, AD may manifest more unilaterally. However, it’s also possible that even absent any true underlying lateralization, one side may simply be more predictive for some patients.

### Conclusion

In this study, we propose an AD diagnosis framework based on deep 3D CNN model using structural MRI, empowered with the inclusion of longitudinal scans. The proposed framework demonstrates high classification performance in Alzheimer’s disease versus cognitive normal. In addition, we demonstrate high accuracy in MCI progression prediction applying the model trained on AD vs. CN classification to the MCI subgroup. Furthermore, through class activation map and rigorous ablation analyses on both slice-level and lobe-level, we pinpoint hippocampal formation as the most predictive regions for AD classification, affirming the prominence of hippocampal formation in AD diagnosis, and demonstrating the importance of regionality even in highly complicated deep neural network models. And importantly, the proposed classification and regional analyses methods constitute a general framework that can easily be applied to other disorders and imaging modalities.

## Acknowledgments

Data collection and sharing for this project was funded by the Alzheimer’s Disease Neuroimaging Initiative (ADNI) (National Institutes of Health Grant U01 AG024904) and DOD ADNI (Department of Defense award number W81XWH-12-2-0012). ADNI is funded by the National Institute on Aging, the National Institute of Biomedical Imaging and Bioengineering, and through generous contributions from the following: AbbVie, Alzheimer’s Association; Alzheimer’s Drug Discovery Foundation; Araclon Biotech; BioClinica, Inc.; Biogen; Bristol-Myers Squibb Company; CereSpir, Inc.; Cogstate; Eisai Inc.; Elan Pharmaceuticals, Inc.; Eli Lilly and Company; EuroImmun; F. Hoffmann-La Roche Ltd and its affiliated company Genentech, Inc.; Fujirebio; GE Healthcare; IXICO Ltd.; Janssen Alzheimer Immunotherapy Research & Development, LLC.; Johnson & Johnson Pharmaceutical Research & Development LLC.; Lumosity; Lundbeck; Merck & Co., Inc.; Meso Scale Diagnostics, LLC.; NeuroRx Research; Neurotrack Technologies; Novartis Pharmaceuticals Corporation; Pfizer Inc.; Piramal Imaging; Servier; Takeda Pharmaceutical Company; and Transition Therapeutics. The Canadian Institutes of Health Research is providing funds to support ADNI clinical sites in Canada. Private sector contributions are facilitated by the Foundation for the National Institutes of Health (www.fnih.org). The grantee organization is the Northern California Institute for Research and Education, and the study is coordinated by the Alzheimer’s Therapeutic Research Institute at the University of Southern California. ADNI data are disseminated by the Laboratory for Neuro Imaging at the University of Southern California.

Data were provided in part by the Human Connectome Project, WU-Minn Consortium (Principal Investigators: David Van Essen and Kamil Ugurbil; 1U54MH091657) funded by the 16 NIH Institutes and Centers that support the NIH Blueprint for Neuroscience Research; and by the McDonnell Center for Systems Neuroscience at Washington University.

## Author contributions statement

XF, SAS, FAP designed the study. XF organized and pre-processed the MRI data. XF proposed and implemented the classification algorithms. XF and JY proposed and implemented saliency based and lobe based regional analysis. XF, ZCL, and FAP proposed and implemented the slice based ablation analysis. All authors wrote the manuscript, with special mention of the machine learning perspective from ZCL.

## Additional information

**Figure S1.**
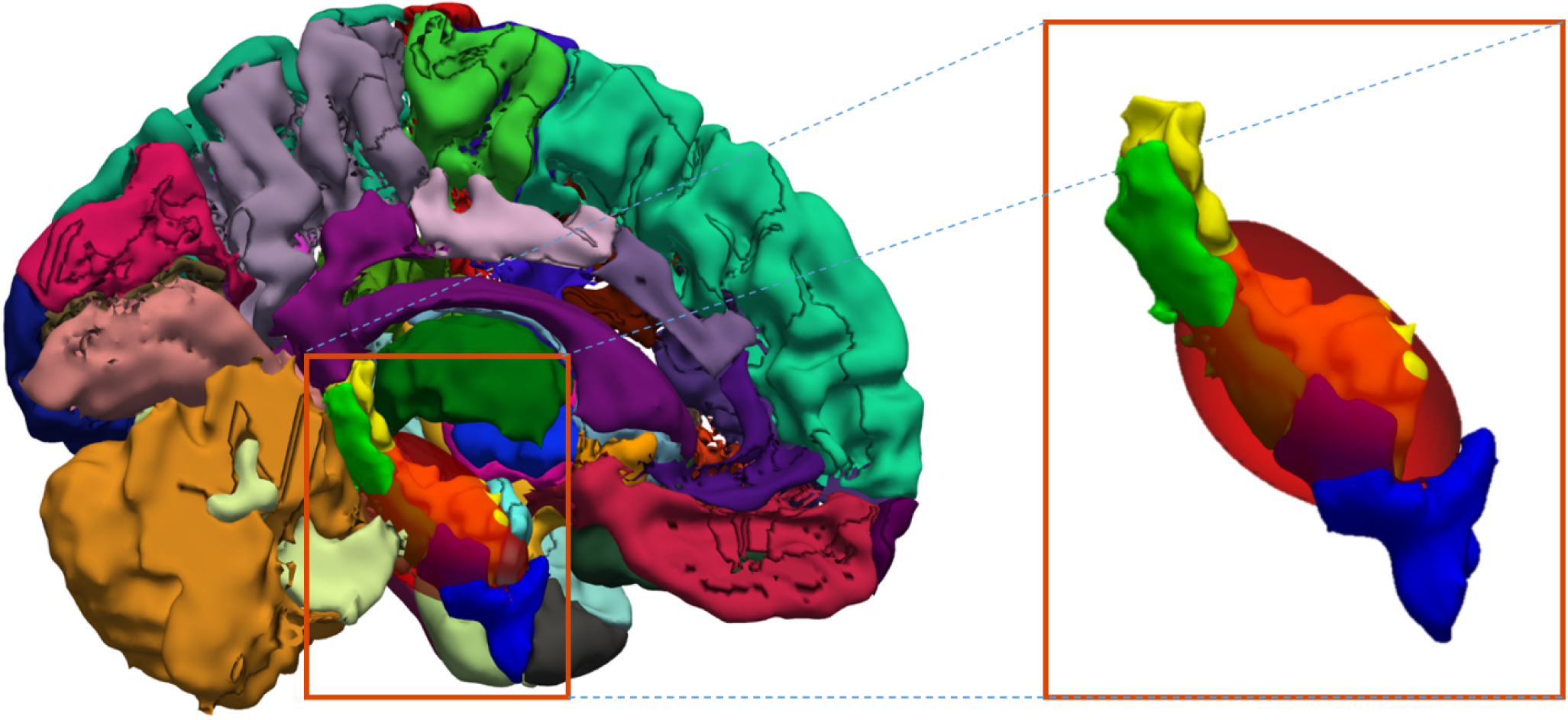
3D rendering of the class activation map (iso-surface at 0.8, red) with the label of hippocampus (yellow), entorhinal cortex (blue), parahippocampal cortex (green) from FreeSurfer template.

**Figure S2.**
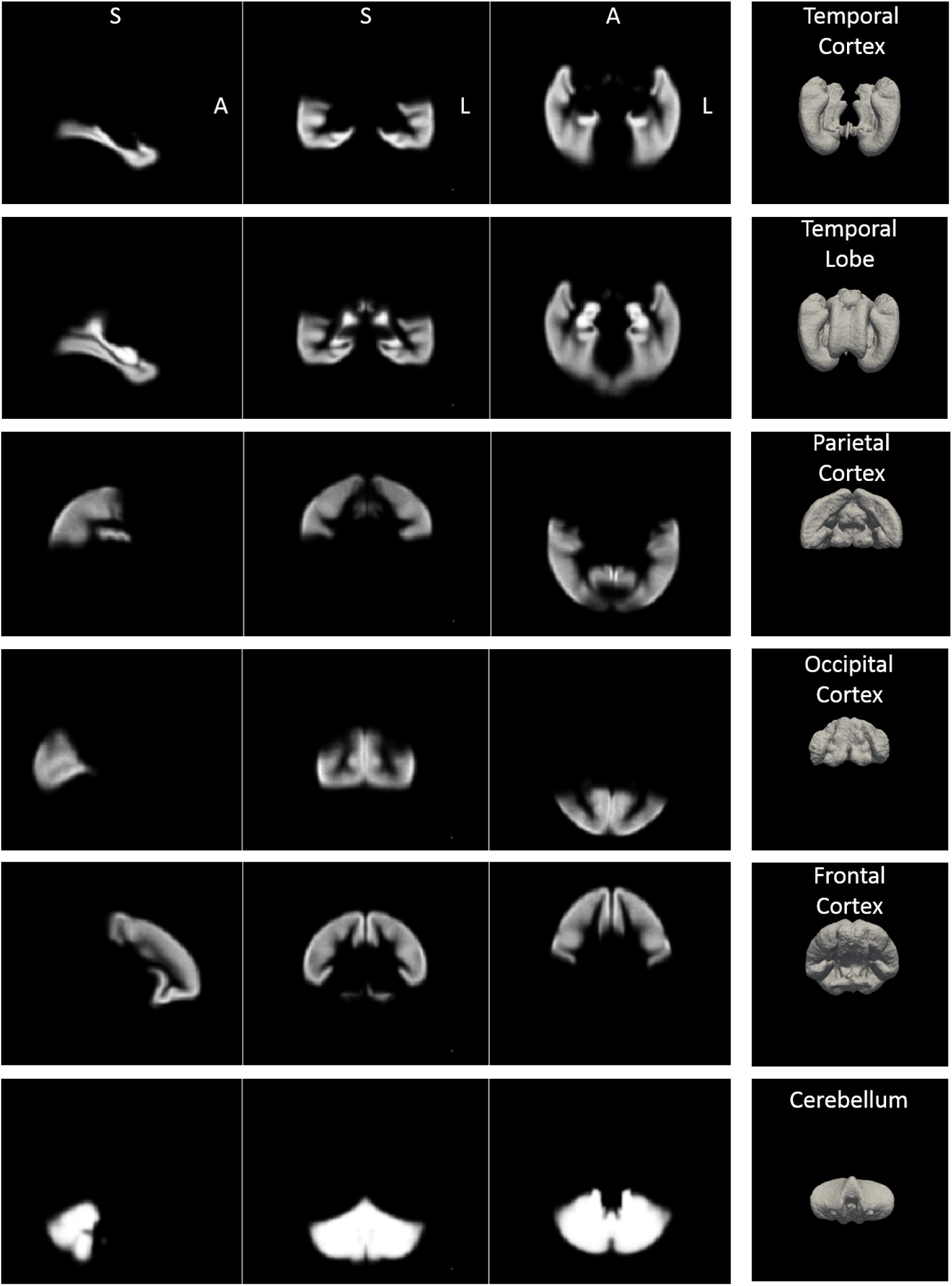
Lobe (and cerebellum) probability maps.

**Figure S3.**
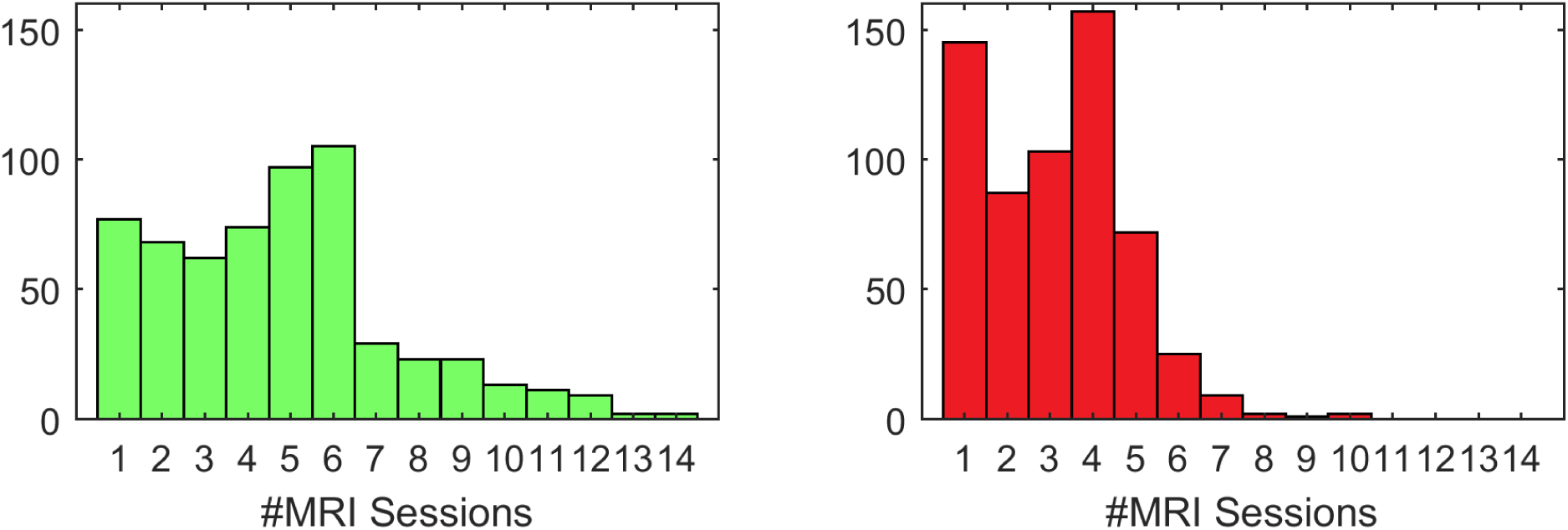
MRI sessions per subject (left) scans of cognitively normal subjects, (right) scans of AD subjects.

## Footnotes

1 http://adni.loni.usc.edu/

2 http://adni.loni.usc.edu/methods/documents/mri-protocols/

3 http://adni.loni.usc.edu/methods/mri-analysis/mri-pre-processing/

4 https://humanconnectome.org/

5 https://surfer.nmr.mgh.harvard.edu/fswiki/CorticalParcellation/

